# A user guide to environmental protistology: primers, metabarcoding, sequencing, and analyses

**DOI:** 10.1101/850610

**Authors:** Stefan Geisen, Daniel Vaulot, Frédéric Mahé, Enrique Lara, Colomban de Vargas, David Bass

## Abstract

Protists – all eukaryotes besides fungi, animals, and plants - represent a major part of the taxonomic and functional diversity of eukaryotic life on the planet and drive many ecosystem processes. However, knowledge of protist communities and their diversity lags behind that of most other groups of organisms, largely due to methodological constraints. While protist communities differ markedly between habitats and biomes, they can be studied in very similar ways. Here we provide a guide to current molecular approaches used for studying protist diversity, with a particular focus on amplicon-based high-throughput sequencing (metabarcoding). We highlight that the choice of suitable primers artificially alters community profiles observed in metabarcoding studies. While there are no true ‘universal’ primers to target all protist taxa as a whole, we identify some primer combinations with a wide taxonomic coverage and provide detailed information on their properties. Although environmental protistan ecological research will probably shift towards PCR-free metagenomics or/and transcriptomic approaches in a near future, metabarcoding will remain the method of choice for in-depth community analyses and taxon inventories in biodiversity surveys and ecological studies, due its great cost-efficiency, sensitivity, and throughput. In this paper we provide a guide for scientists from a broad range of disciplines to implement protists in their ecological analyses.

## Introduction

The past three decades have seen massive developments in our understanding of microbial diversity in most of the Earth ecosystems. Molecular approaches, particularly the advent of high-throughput sequencing (HTS), have enabled cultivation-independent diversity analyses of microorganisms. PCR-based amplicon HTS (here defined as metabarcoding) data for bacteria and archaea have rapidly accumulated throughout this period, revealing a vast and mostly unknown taxonomic diversity (Gans, Wolinsky, & Dunbar, 2005; Leininger et al., 2006; Louca, Mazel, Doebeli, & Parfrey, 2019; Parks et al., 2017; Sogin et al., 2006). While sequencing errors were shown to have artefactually inflated the inferred diversity in many early studies (Huse, Huber, Morrison, Sogin, & Welch, 2007), increased quality control and refined techniques have subsequently confirmed that the global diversity of prokaryotes likely exceeds a million bacterial species-level lineages (Louca et al., 2019; Parks et al., 2017; Thompson et al., 2017), with many previously unknown clades emerging. Similarly, fungal diversity has been studied extensively using HTS and taxon diversity has been estimated to exceed 100,000 (Buée et al., 2009; Tedersoo et al., 2014), if not an order of magnitude greater (Bass & Richards, 2011; Hawksworth & Lucking, 2017). Protist metabarcoding is lagging behind that of prokaryotes and fungi (Geisen et al., 2017), yet suggesting global species-level diversity estimates in the millions (de Vargas et al., 2015; Mahé et al., 2017).

Protists are of key importance for ecosystem functioning and are major drivers in diverse nutrient cycling pathways. Autotrophs (algae) are the counterparts of land plants as main carbon fixers in aquatic environments (Worden et al., 2015). Heterotrophic protists catalyse nutrient cycling in aquatic and terrestrial environments as selective consumers of bacteria and fungi, being critical drivers of the so-called ‘microbial loop’ (Azam et al., 1983; Bonkowski, 2004). Parasitic protists influence community dynamics of larger eukaryotic hosts, including plants and animals (S. Geisen et al., 2018; Mahé et al., 2017; Worden et al., 2015). Through these abiotic and biotic processes, protists make nutrients available to either smaller or larger members of food webs and enhance eco-system dynamics (Bonkowski, 2004) and/or connectivity (Lima-Mendez et al., 2015). Nevertheless, the diversity of protists is often excluded from microbiome and food-web studies (Lundberg et al., 2012; Mendes et al., 2011). Therefore, we are missing information of an important biotic component that regulates ecosystem functioning.

The protist biodiversity knowledge gap at least partly originates from methodological constraints. Standardized PCR-based approaches with a relatively limited number of recommended primer pairs are applied to study bacteria (J. Gregory Caporaso et al., 2010; J. G. Caporaso et al., 2011; Cole et al., 2014; Knight et al., 2018; Walters et al., 2016) and fungi (Ihrmark et al., 2012; Lindahl Björn et al., 2013; Nilsson et al., 2019). However, a consensus on comparable molecular approaches using a standardized PCR-based approach to study protists does not exist. Several primer sets are regularly used (Adl et al., 2019; Adl, Habura, & Eglit, 2014; Hadziavdic et al., 2014), although their benefits and shortcomings are generally poorly known. Ideally, a PCR protocol would amplify all members of the target group with equal likelihood, avoid excessive amplification of non-target lineages, and provide amplicons with high taxonomic resolution. The target sequence should also be well represented in public databases, and provide enough phylogenetic signal for eco-evolutionary analyses. However, a single such ideal PCR system cannot be developed. In reality, each primer set has some phylogenetic bias, and the use of many different sets therefore limits comparability across studies (Fouhy, Clooney, Stanton, Claesson, & Cotter, 2016; Ramirez et al., 2018; Stoeck et al., 2010). In fact, primer pairs have often been designed to answer specific questions in protist community profiling (see below), based on the experience or preference of individual researchers. A comparative synthesis of existing ‘protist primers’ for non-experts is still lacking. We therefore review some of the most frequently used primer pairs, evaluate their pros and cons, and provide suggestions for optimal primer choice in different environmental settings. We then introduce guidelines to consider in HTS-based analyses of protist biodiversity. Lastly, we provide an overview of future opportunities and challenges introduced by PCR free HTS approaches.

### 18S rRNA gene: the universal marker to assess protist biodiversity

The 16S (small subunit) ribosomal RNA gene (Woese, 1987) has been used as a genetic marker for prokaryotic (bacteria and archaea) diversity since the earliest days of amplicon-based diversity studies (Giovannoni & Cary, 1993; Pace, 1997). Its eukaryotic counterpart (18S) was later adopted for studies of micro-eukaryotes (Lopez-Garcia, Rodriguez-Valera, Pedros-Alio, & Moreira, 2001; Moon-van der Staay, De Wachter, & Vaulot, 2001; Moreira & López-Garcia, 2002; van Hannen et al., 1999). The 16S/18S rRNA gene encodes the RNA molecule that forms part of the small subunit of the ribosome. The ribosome is a macro-molecule essential to all organisms to translate mRNA into proteins, its function thus being conserved across the tree of life. While other markers are sometimes preferable for finer-resolution, often group-specific, biodiversity assessment (Fig. 2 in Jan Pawlowski et al. (2012)), the 18S rRNA is the best suited gene for exploring the diversity of protists as a whole. The 18S rRNA gene contains nine relatively variable regions flanked by more conserved regions that together span a range of evolutionary rates, enabling phylogenetic comparisons of both distant and closely related taxa (Neefs, Van de Peer, De Rijk, Chapelle, & De Wachter, 1993). Primers can be designed in more or less conserved regions to target a wide diversity of taxa. However, there are important considerations to be made when choosing or designing primer sets to amplify 18S rRNA gene regions. Despite being most comprehensive, NCBI’s GenBank, is not recommendable to classify 18S rRNA gene sequences due to large amount of errors that include wrongly annotated sequences. At least two general databases are dedicated to the 18S rRNA gene, Silva (Pruesse et al., 2007) and PR^2^ (Guillou et al., 2013), while more specialized databases have been designed for specific taxonomic groups, e.g. foraminifera (Morard et al., 2015) or dinoflagellates (Mordret et al., 2018).

**Fig. 1.**
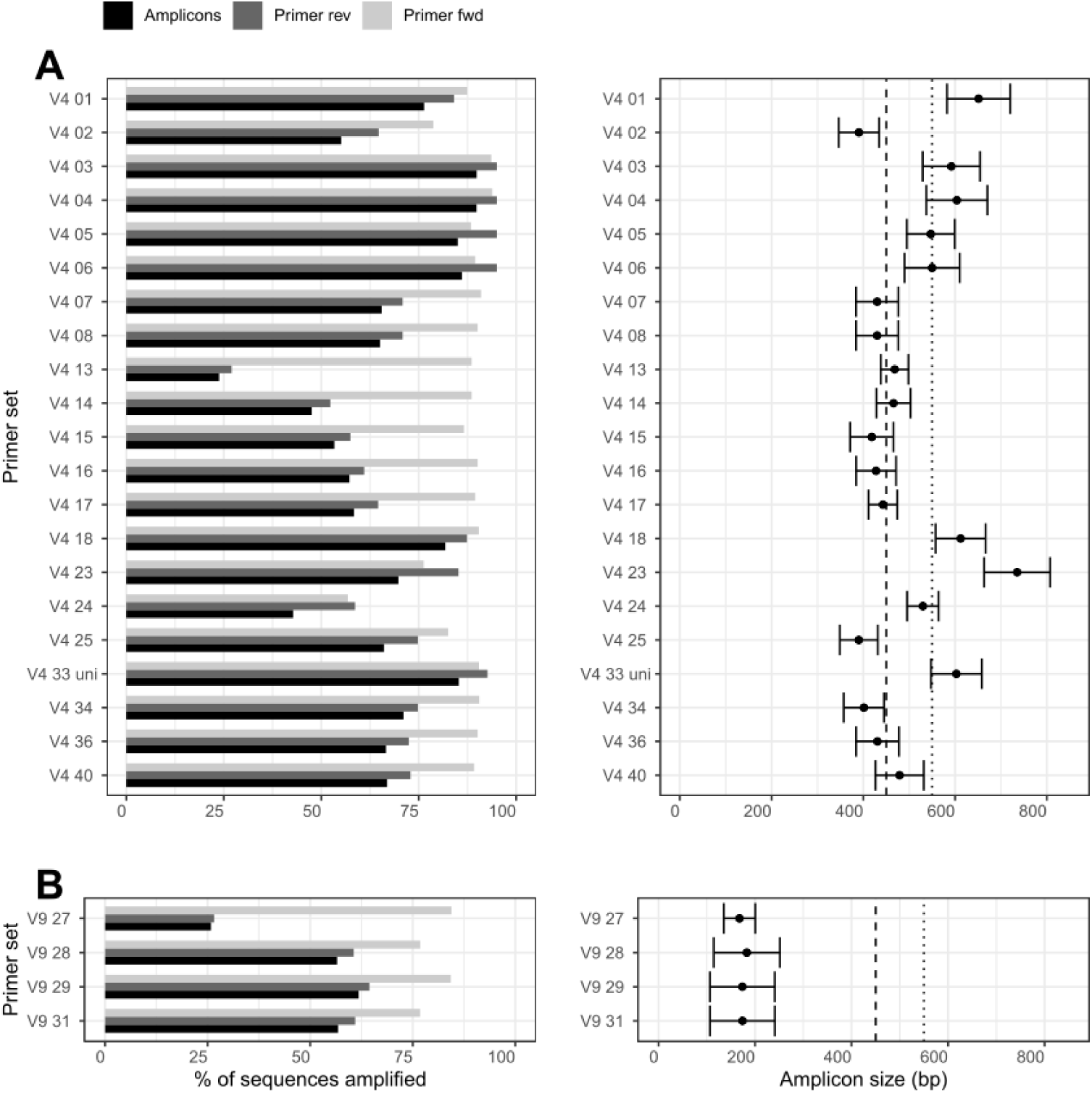
Left panel. Percentage of reference sequences with a perfect match to both forward and reverse primers individually and the entire (A) V4 and (B) V9 18S rRNA gene region; right panels show amplicon sizes targeted by different primer pairs including lengths that can be covered by the most commonly used Illumina sequencers (dotted line: 2×300 base pairs; dashed line: 2×250 base pairs); error bars represent the standard deviation. PR^2^ version 4.11.1 was used the reference 18S rRNA database. Note that this figure provides an overview of all currently used primer sets to target protists; details for each of the primers is given in Table 1.

**Fig. 2.**
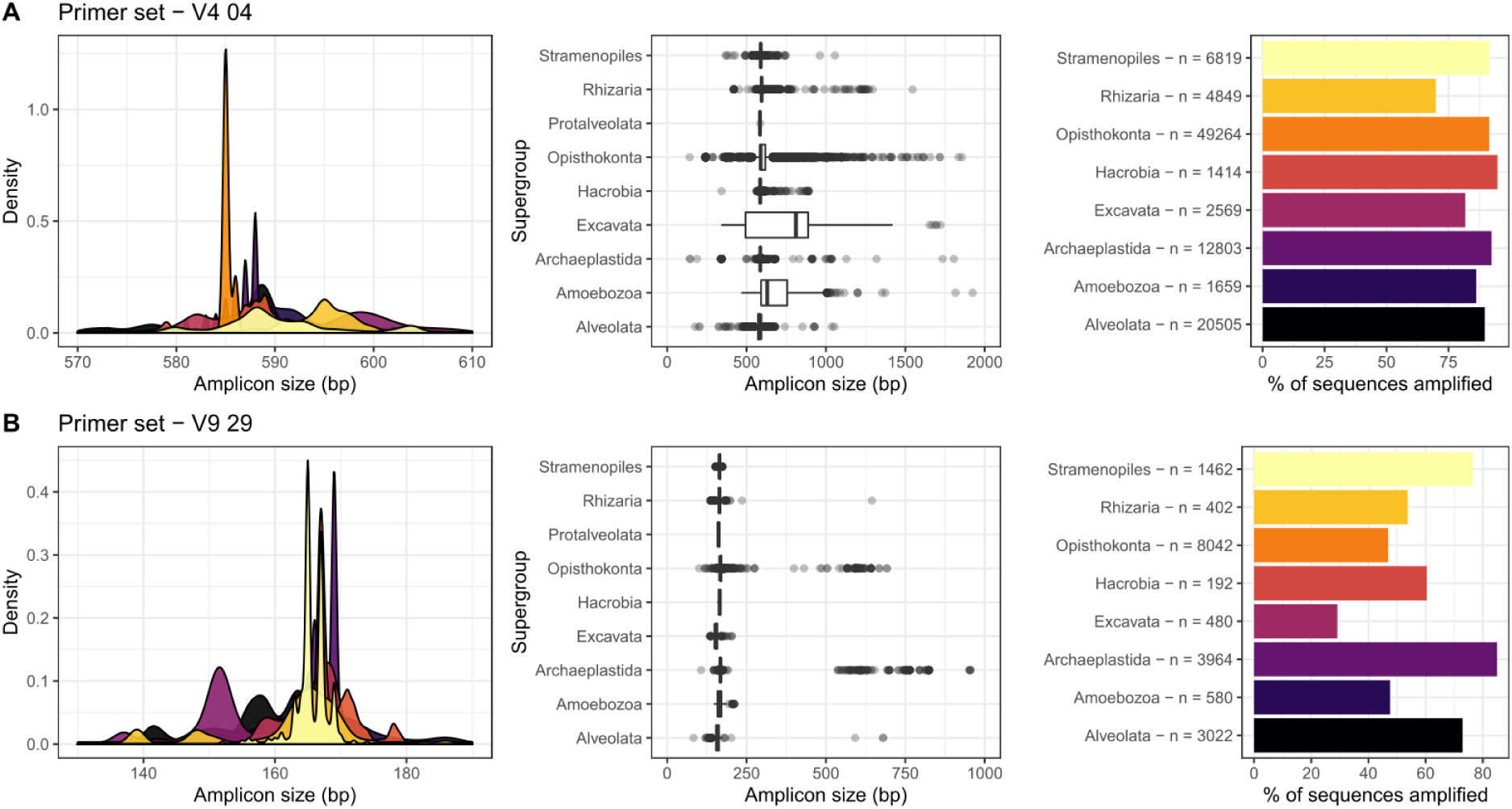
Amplicon lengths differences of higher taxonomic level protist lineages exemplified for each of a broadly targeted V4 (A) and V9 (B) primer set (Table 1). Lengths differences are prevalent between protist groups especially in the V4 region leading to differential amplification efficiency between the groups. Note that Hacrobia represents the sum of Haptophytes, Cryptophytes and Centrohelids. n represents the number of taxa for a given supergroup present in PR^2^.

### A comparative overview of 18S rRNA gene primers with large taxonomic spectrum

HTS protist metabarcoding studies have employed dozens of different primers in various combinations, targeting a range of variable 18S rRNA gene regions. Among the most widely targeted regions in early protist-focused HTS metabarcoding studies was the V1-3 region (Euringer & Lueders, 2008), a region with relatively high taxonomic resolution (Hadziavdic et al., 2014; Tanabe Akifumi et al., 2015). However, this region is decreasingly being used to target protists due to insufficient reference sequence coverage of the 5’ end of the 18S rRNA gene, its excessive length for most current HTS methods, and unsuitable regions for primer annealing (Hugerth et al., 2014). Instead, the V4 and V9 regions are most commonly used nowadays; they are the focus of this review, as summarised in Table 1.

**Table 1.**
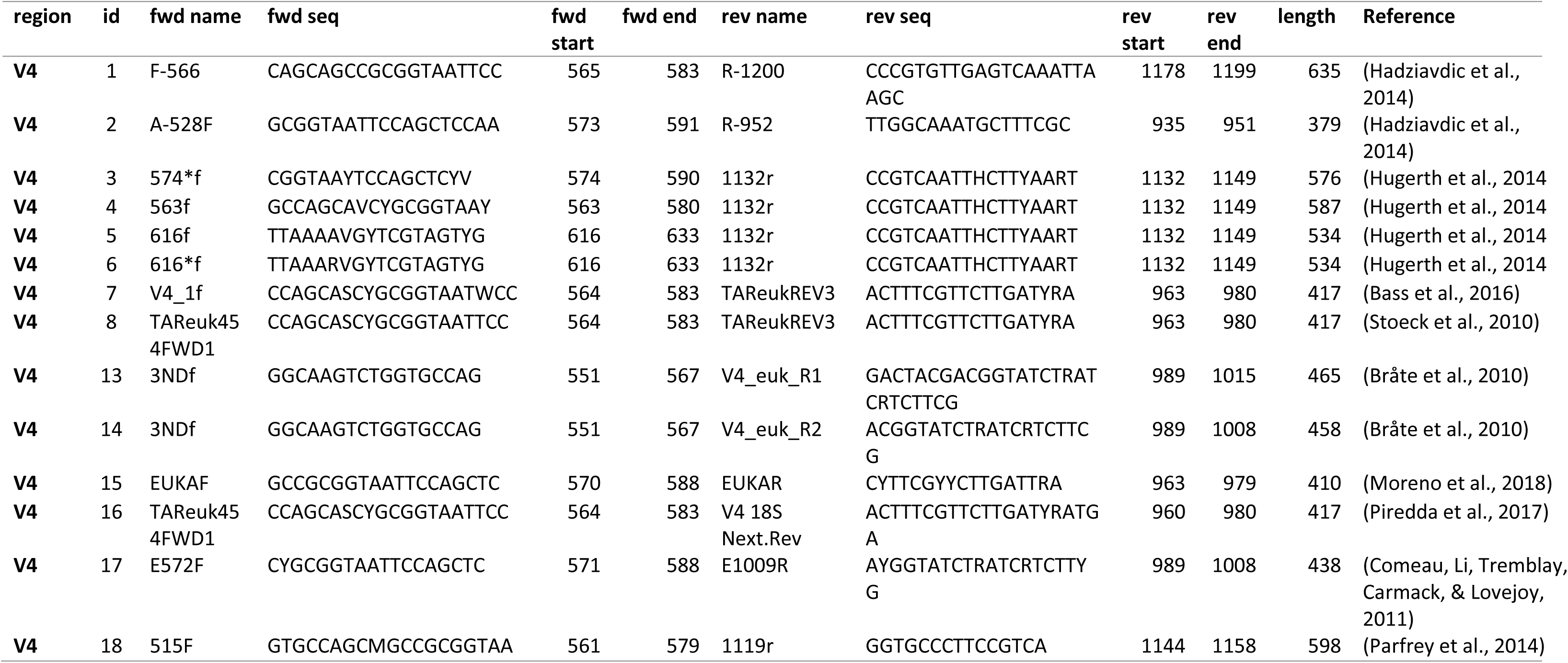
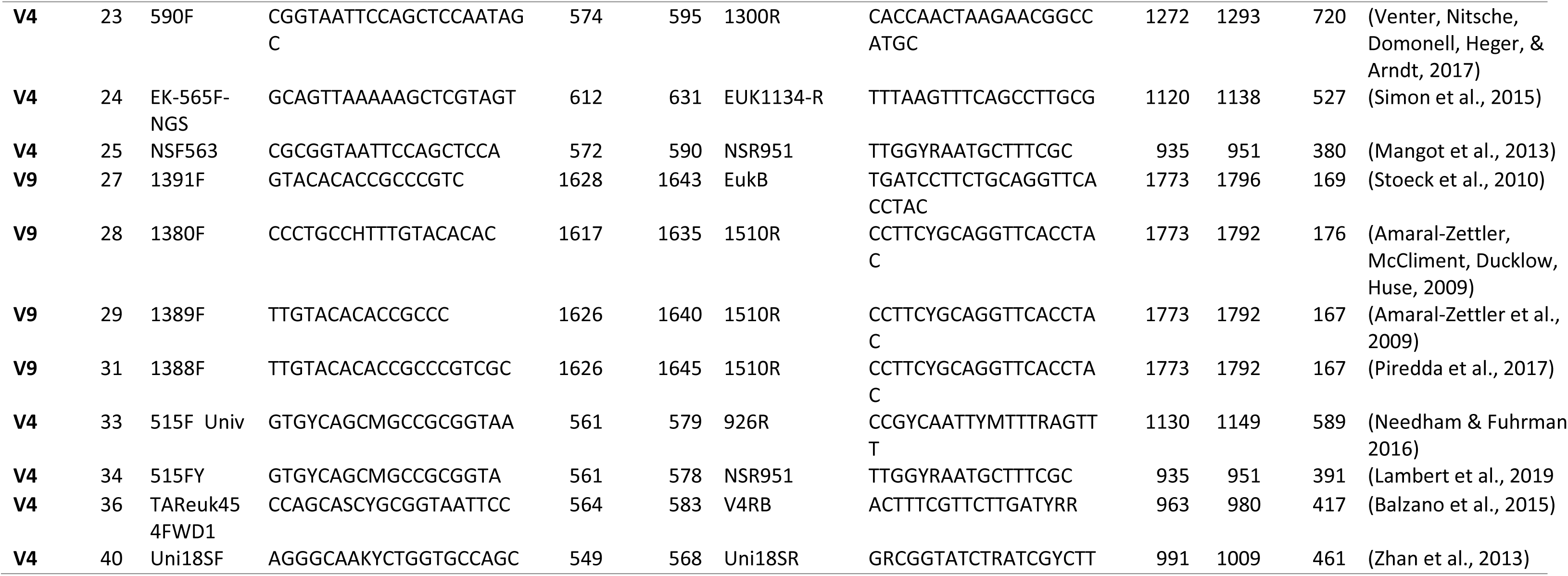
Primer sets targeting the V4 and the V9 region that were used in metabarcoding high-throughput sequencing efforts to study protists; id: primer pair identifier; #: forward or reverse primer name; Fwd: forward primer; Rev: reverse primer; fwd and rev start/end: positions on *Saccharomyces cerevisiae* sequence FU970071; note: id33 also targets prokaryotes; id35 selects against metazoan sequences.

The V9 region was the most frequently used metabarcoding target in early high throughput studies as its relatively short amplicon length (< 200 bp) was better suited to the sequence length limitations of early HTS technologies. This short length makes the V9 region a suitable target also for contemporary ultra-deep metabarcoding technologies, making it a most cost-effective approach to study protist community diversity, while allowing stringent quality control. The downsides of the V9 region is associated with its short length (Fig. 1), providing limited phylogenetic information. As such, the 400-500 bp V4 region fragment is increasingly used (Table 1). Furthermore, the V4 region is closest of all HTS-suitable 18S rRNA gene regions to that of the full-length gene and allows the high phylogenetic/taxonomic resolution often to species- or genus-level (Dunthorn, Klier, Bunge, & Stoeck, 2012; Hu Sarah et al., 2015).

The V4 region also enables the most accurate taxonomic placement of unassigned HTS amplicon sequences (Mahé et al., 2017). For these reasons it has been suggested as the main universal protist barcode region (Jan Pawlowski et al., 2012). However, some disadvantages are associated with the V4 region such as its length, which is too long for some short-read optimized sequencing methodologies such as Illumina HiSeq and NovaSeq (Fig. 1, Fig. 2). V4-based metabarcoding surveys also suffer from PCR/sequencing biases due to amplicon length variability (Fig. 1) and variable secondary structures.

### Challenges of 18S rRNA gene targeted diversity analyses

The 18S rRNA gene is more complex than the prokaryotic 16S rRNA gene. It is on average three hundred base pairs longer (appr. 1,800 bp) and displays profound evolutionary rate variation between different taxa. This renders the design of truly universal primers impossible (Adl et al., 2014; Hadziavdic et al., 2014). Primers described as ‘universal’ are better thought of as ‘broadly-targeted’. Furthermore, even the most broadly-targeted primer pair will not amplify all protist lineages equally well, such as higher taxonomic ranks (e.g. eukaryotic supergroup; Fig. 3), but differential amplification exists also at lower taxonomic ranks such as at the class level (Fig. 4). Some highly divergent lineages will not amplify at all, as their 18S rRNA gene sequences can differ in multiple nucleotide positions even in the most conserved primer sites. This is the case for many parasitic protists (Bass, Stentiford, Littlewood, & Hartikainen, 2015; Hartikainen et al., 2014), but also diverse free-living lineages (Bass et al., 2016; Bass, Yabuki, Santini, Romac, & Berney, 2012; Fiore-Donno, Weinert, Wubet, & Bonkowski, 2016); at high taxonomic rank this is most obvious for Discoba (within Excavata; Fig. 2). For those groups, alternative approaches are needed to study their communities, such as applying group-specific primers. Even when primers are a perfect sequence match to the template, differences in length and secondary or tertiary structures of the amplified region, due to insertion-deletions (indels) or the presence of introns, which leads to biases, as shorter and structurally simpler fragments are preferentially amplified over longer ones (Figs. 2 and 3). This can influence the sequence representation of protists at all levels, even at supergroups. For instance, Stramenopiles and Alveolates have relatively short V4 amplicon lengths leading to preferential PCR-amplification, while the longer V4 regions of Discoba (including Excavata) and Amoebozoa (Figs. 2 and 3) amplify less efficiently (Geisen, Laros, Vizcaíno, Bonkowski, & de Groot, 2015).

**Fig. 3.**
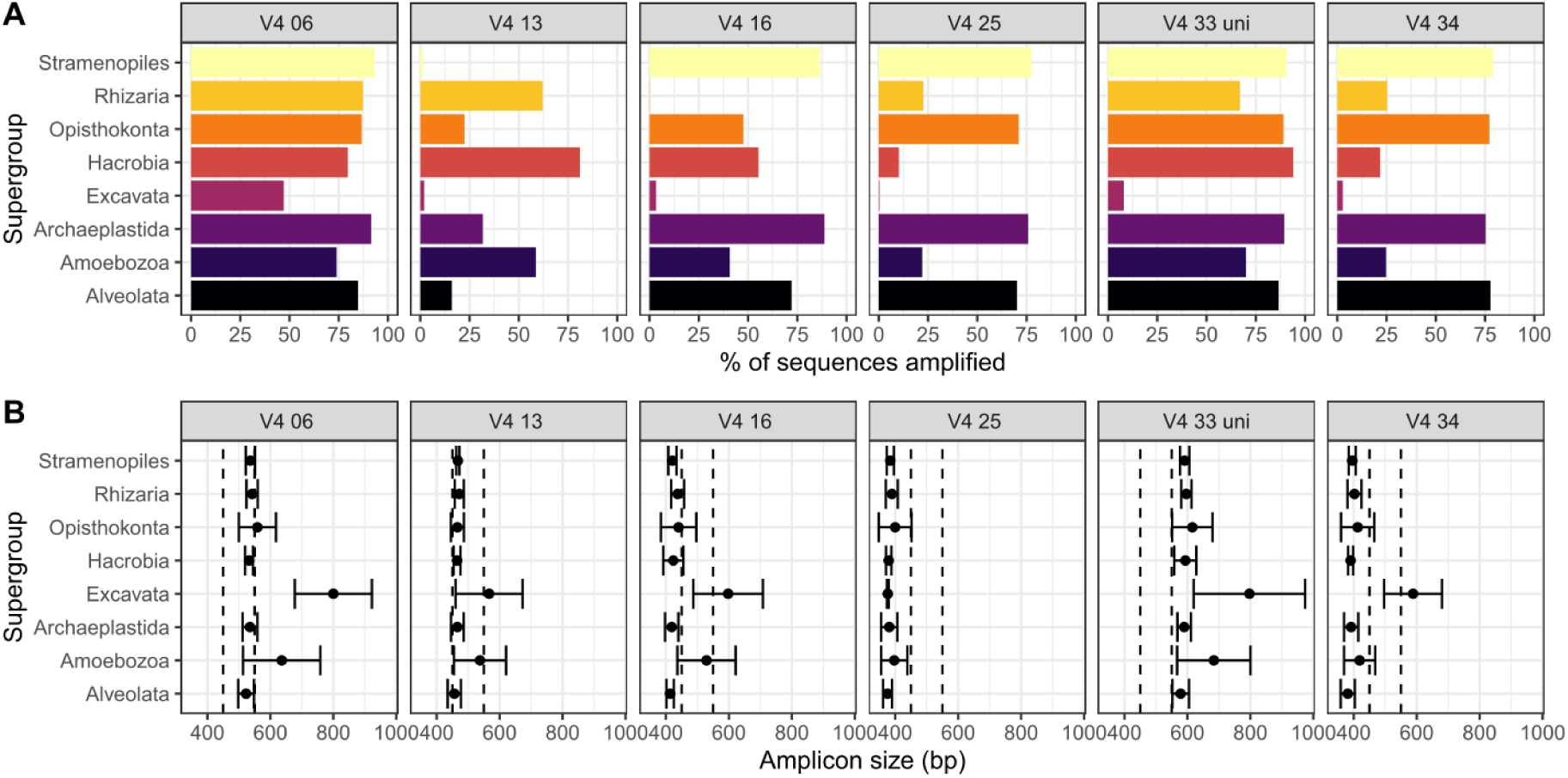
Coverage of some of the most broadly targeting primer pairs (Fig. 1, Table 1) as identified with perfect matches of both primer pairs to the target sequences for the main protist lineages. This shows that (A) primers do not equally amplify higher taxonomic level lineages of protists and (B) amplicon lengths differ between supergroups and depending on primer sets. Note that Hacrobia represents the sum of Haptophytes, Cryptophytes and Centrohelids. For details on all primer sets see Suppl. Fig. 1.

**Fig. 4.**
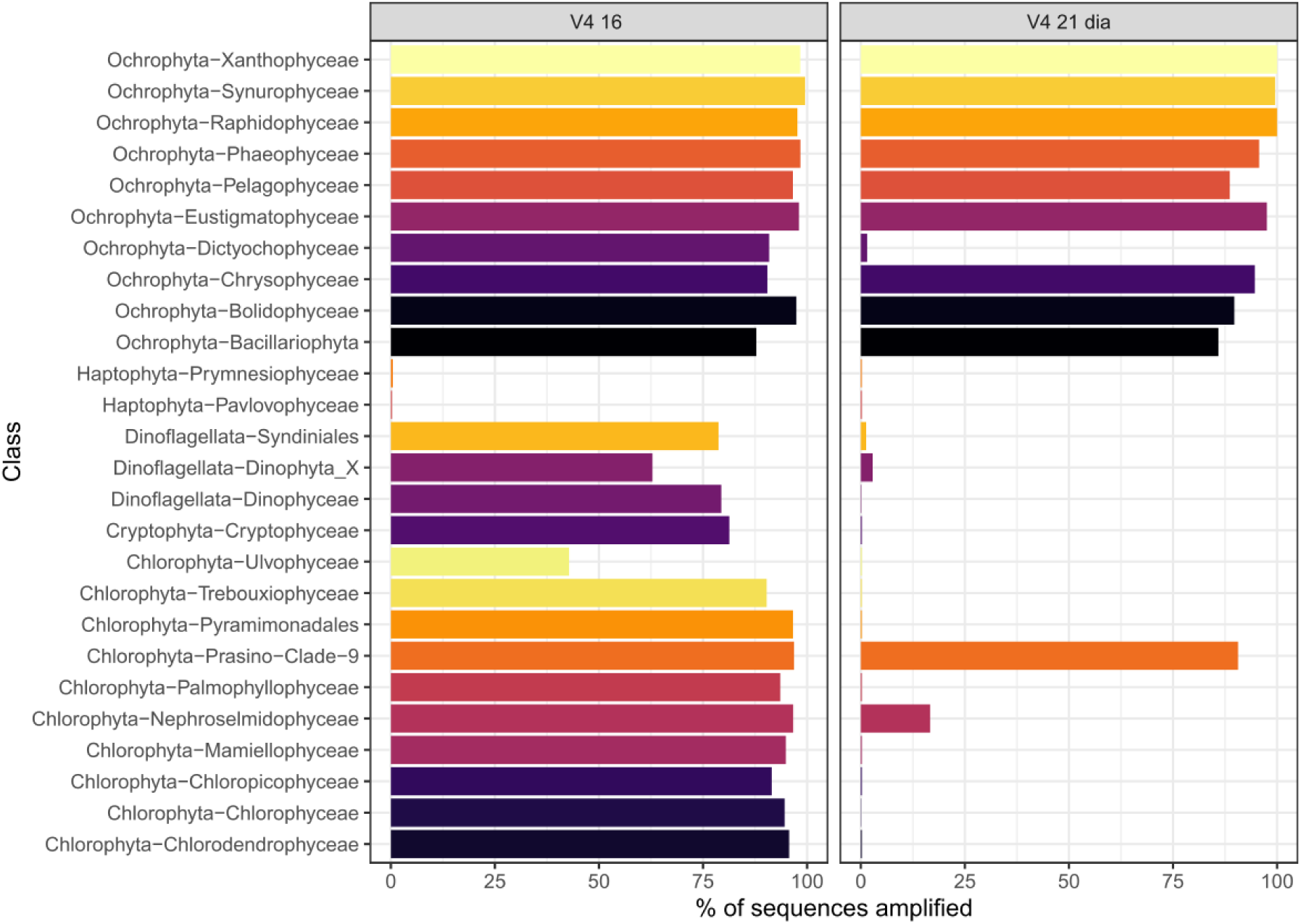
Coverage differences at higher taxonomic resolution (which corresponds, approximatively, to Class level) as identified with perfect matches of both forward and reverse primers illustrated for primer sets 16 and 21, showing that amplification success differences between protist groups (see Fig. 2) can also be present at lower taxonomic levels. The “universal” primer set 16 does not amplify Haptophyta *in silico*, but in fact partially amplified them in natural communities *in vivo* likely because the mismatch is located in the middle of the primer (Piredda et al., 2017). Primer set 21 which is described as specific of diatoms (Zimmermann, Jahn, & Gemeinholzer, 2011) also amplifies other Ochrophyta as well as some green algae.

The ribosomal gene operon is present in multiple copies, and copy numbers vary greatly between taxonomic groups by up to three orders of magnitude (Gong, Dong, Liu, & Massana, 2013; Zhu, Massana, Not, Marie, & Vaulot, 2005). There is evidence in some protist groups that the number of 18S rRNA gene reads (suggesting gene copies) positively correlates with relative cell abundance (Giner et al., 2016), biomass (Pitsch et al., 2019) or biovolume (de Vargas et al., 2015), but these correlations do not hold across divergent protist groups (Pitsch et al., 2019). Variable copy numbers between lineages can profoundly alter interpretation of relative abundance or biomass of taxa within protist communities in metabarcoding studies. Inferred lineage diversity can also be biased by intragenomic variation between rRNA gene copies. The degree of intragenomic variation of rRNA gene copies is generally low (0-1%), although in some groups it can reach several percent in some ciliates (Gong et al., 2013), Myxomycetes (García-Martín, Zamora, & Lado, 2019), and Foraminifera (Weber & Pawlowski, 2014). Both of these factors can vary greatly between closely related taxa (Andre et al., 2014; García-Martín et al., 2019; Gong et al., 2013; Nassonova, Smirnov, Fahrni, & Pawlowski, 2010; Weber & Pawlowski, 2014). This can artificially increase diversity estimates as single taxa can then be treated as different species or, if intraspecific sequence diversity is high, as different taxonomic groups (Caron & Hu, 2019).

The phylogenetic resolution a barcoding region provides is key for HTS studies. However, because taxonomic marker gene evolution occurs at different rates relative to phenotypic evolution across protist groups, there is no fixed or consensus relationship between marker genes dissimilarity and taxonomy (Boenigk, Ereshefsky, Hoef-Emden, Mallet, & Bass, 2012). Therefore, fixed percentage differences between lineages cannot be used to infer taxonomic distances between groups, and often within them. Different solutions to this can be approached for individual groups, such as targeting a combination of diverse variable regions or using barcoding regions other than the 18S rRNA gene. This, however, is often not feasible in larger-scaled ecological studies due to cost-limitations, and a lower resolution has to be accepted. Another approach is the use of selective targeted bioinformatics analysis for specific groups of interest, for example mapping short reads onto robust phylogenetic trees constructed with longer sequences.

Lastly, a metabarcoding approach targeting protists will depict a community of protists that differs from the protist community present *in situ* or *in vivo*, because of the above-mentioned taxon-specific PCR biases and copy number differences. As such, the primer pairs chosen alter the inferred composition of protist communities in a primer-pair specific manner, which is illustrated in studies applying primers targeting both the V4 and V9 regions (Pagenkopp Lohan, Fleischer, Carney, Holzer, & Ruiz, 2016; Stoeck et al., 2010; Tragin, Zingone, & Vaulot, 2018). For example, V9 datasets generally contain relatively more sequences that can only be assigned at the domain level (eukaryotes) compared to V4 assignments (Pagenkopp Lohan et al., 2016).

### Group-specific primer sets

Lineages that do not amplify well with broadly-targeted primers or when non-target sequences are expected to dominate the results can be targeted in metabarcoding approaches with group-specific primers. These target a range of different 18S rRNA gene regions (Supplementary Table 1). For instance, Cercozoa have specifically been targeted in soils to avoid amplification of fungi (Fiore-Donno Anna et al., 2017; Harder et al., 2016; Lentendu et al., 2014). However even within Cercozoa, amplification of some clades requires even more specific primers, such as the divergent coprophilic *Helkesimastix*-*Guttulinopsis*-*Rosculus* clade (Bass et al., 2016). Supplementary Table 1 provides a summary of group-specific primer sets that have so far been used in 18S V4 region rRNA gene protistan metabarcoding efforts. Note that this table only includes primers that have been used and proven successful in HTS metabarcoding efforts. Many other primer sets have been proposed (Adl et al., 2019; Adl et al., 2014) that might be applicable but have not yet been tested in metabarcoding approaches.

Genes other than the 18S rRNA can be targeted to increase the taxonomic resolution of specific protist lineages(Jan Pawlowski et al., 2012). For instance, ITS sequences have been used to assess the distribution of uncultivated heterotrophic marine protists (Rodríguez-Martínez, Rocap, Logares, Romac, & Massana, 2011), often providing species-level resolution(Stern et al., 2012). The first subunit of the mitochondrial cytochrome oxidase as the universal barcoding region in animals (Hebert, Ratnasingham, & deWaard, 2003) also allows high-taxonomic discrimination of many protist taxa (Heger et al., 2011; Singer et al., 2018). The binding sites of these primers are so conserved that the same protocols as for animals have been used for several divergent Amoebozoa (Kosakyan et al., 2012; Nassonova et al., 2010). For specific groups of photosynthetic protists such as diatoms the *rbcL* gene can be used (Rimet et al., 2019).

An alternative to group-specific primers that target specific groups of protists is the use of primer combinations that amplify broadly but exclude certain groups of protists and other eukaryotes. As such, fungal sequences can be avoided in soils (Fiore-Donno et al., 2016; Geisen, 2016; Lentendu et al., 2014), and in holobionts for the exclusion of plant or metazoan host-associated sequences.

### Assessing host-associated protist diversity

The highly parallel throughput of currently available sequencing techniques means that protist diversity can be effectively sampled and analysed even in samples dominated by non-protist taxa. However, when targeting host-associated protists, co-amplification of host DNA can overwhelm the protist signal. In these cases, group-specific primer sets can be used to focus on a particular group. This strategy is typically used for high-resolution diversity analyses of fungi or Peronosporomycetes (formerly Oomycetes) inside plant roots (Ramirez et al., 2019; Sapkota & Nicolaisen, 2015). However, the disadvantage of this approach is that multiple groups that might be of ecological importance as symbionts cannot be simultaneously targeted. To fully assess the eukaryotic microbiome (‘eukaryome’ (Javier del Campo, Bass, & Keeling, 2019)) in animal hosts, broadly-targeted eukaryotic 18S rRNA gene V4 region primers that avoid amplification of most animal sequences are available (Bower et al., 2004; Javier del Campo et al., 2019). Alternatively, blocking primers can also be applied that reduce amplification of selected, non-target DNA and thereby increase the amount of target DNA (Tan & Liu, 2018; Vestheim & Jarman, 2008). To the best of our knowledge, plant-blocking primers have not yet been developed to enable amplification of protists within or on plant tissue.

#### Downstream sequence analyses

Downstream analyses of raw sequence reads are needed process the data and to ensure high data quality. The key steps are common to analyses of all metabarcoding datasets: merging paired-end sequence reads, and quality control steps such as removing low quality, short-read and chimeric sequences (J. Gregory Caporaso et al., 2010). For the subsequent clustering step of processed reads into sequence types representing taxonomic units, an increasingly wide choice of methods is available that significantly impacts measures of diversity and subsequent interpretation. This step aims to eliminate erroneous sequences created during PCR and sequencing, along with sequences originating from intragenomic and intraspecific polymorphisms (Cédric Berney, Fahrni, & Pawlowski, 2004; Richards & Bass, 2005) that can artificially inflate diversity estimates.

Operational Taxonomic Unit (OTU) clustering commonly groups sequences by a user-defined percentage similarity – typically 97 to 99% - using software such as mothur, qiime, usearch, vsearch, etc. (Edgar, 2010; Rognes, Flouri, Nichols, Quince, & Mahé, 2016), using more flexible high taxonomic resolution approaches such as the clustering method SWARM (Mahe, Rognes, Quince, de Vargas, & Dunthorn, 2014, 2015) or the denoising method implemented in dada2 to produce amplicon sequence variants (ASVs). This last approach is now widely used for bacterial communities across the globe (Callahan, McMurdie, & Holmes, 2017; Delgado-Baquerizo et al., 2018; Thompson et al., 2017), and is increasingly used to analyse protist diversity (Chénard et al., 2019). It is possible that the application of ASVs might lead to an over-estimate of protistan diversity as intragenomic gene copy variants or and artefactual sequences may be interpreted as separate taxonomic units (Caron & Hu, 2019; Xiong et al., 2019).

Several methods exist for taxonomic annotation of OTUs/swarms/ASVs: assignment by the Uclust consensus taxonomy assigner, blast, GGSearch (Pearson, 2014), SortMeRNA, RTAX, Kraken or naïve Bayes classification (RDP Classifier (Wang, Garrity, Tiedje, & Cole, 2007)). Two main curated reference databases are available for annotation of 18S rRNA gene reads: SILVA (Pruesse et al., 2007) and PR^2^ (Guillou et al., 2013) (https://github.com/pr2database/pr2database). SILVA incorporates all existing 18S rRNA sequences and relies on automatic annotation, while PR^2^ focuses on the coherence of a taxonomy framework, for example using resources such as Algaebase (Guiry & Guiry, 2008) for photosynthetic protists, and on taxonomic re-annotations performed by supervised pipelines such as EukRef (Boscaro et al., 2018; Javier del Campo et al., 2018). Taxonomic annotation of the same dataset to different databases can result in discrepancies (Dupont, Griffiths, Bell, & Bass, 2016). A community based and expert driven, universal taxonomic framework for protist is underway (UniEuk (C. Berney et al., 2017)) and aims at providing a unified reference taxonomy for protists that will be implemented through the European Bioinformatics Institute (EBI).

### Beyond currently applied metabarcoding approaches

Some new sequencing technologies can generate far longer reads than previously possible (Clarke et al., 2009). Amplicons of thousands of base pairs can be generated with platforms such as Pacific Biosciences and Oxford Nanopore. The potential of these long-read platforms to increase phylogenetic inference and obtain species-level resolution has been shown for fungi and bacteria (Benítez-Páez, Portune, & Sanz, 2016; Tedersoo, Tooming-Klunderud, & Anslan, 2017), and recently for protists (Jamy et al., 2019). Direct RNA sequencing without a cDNA intermediate is also now possible and could allow untargeted protist and general microbiome community profiling (Graham et al., 2019; Marinov, 2017).

In comparison to sequencing platforms focusing on increased read lengths, other platforms (especially Illumina) focus on increased sequencing depth. These are particularly suitable for PCR-free “omics” strategies, including whole community profiling via metagenomics and metatranscriptomics (Carradec et al., 2018; Prosser, 2015). When PCR amplification to increase product yield is avoided, -omics methods circumvent PCR/global-amplification biases that distort diversity analyses and provide community profiles of all three domains of life plus viruses, as well as functional gene information (Flues, Bass, & Bonkowski, 2017; Karsenti et al., 2011; Pesant et al., 2015; Turner et al., 2013; Urich et al., 2008). The systematic collection of multi-omics data (metagenomic, metatranscriptomic, single-cell omics) in aquatic (Carradec et al., 2018; Cuvelier et al., 2010; Seeleuthner et al., 2018) and soil (Geisen, Tveit, et al., 2015; Jacquiod et al., 2016; Turner et al., 2013) systems are unveiling a more true picture of diversity and community composition of protists. A drawback of omics approaches is the high cost per sample since a much deeper sequencing of 10-100 million reads per sample is required. Coverage of any particular target group is also limited, as metagenomic datasets may be dominated by bacterial sequences or host sequences from animal/plant samples (Tedersoo et al., 2015; Urich et al., 2008). The key opportunity for protists afforded by -omics approaches, i.e. the elucidation of patterns of gene expression and interactions, is so far constrained by the deficiency of reference databases compared to those available for prokaryotes (Caron et al., 2017). As this information gap is increasingly filled, however, functional -omics information will help us to better understand ecosystem processes beyond taxonomy-based inferences of protist functions that are currently largely limited to nutrient uptake modes (Stefan Geisen et al., 2018). The first studies linking protist taxa to biological functions using -omics data have revealed the immense potential of these approaches (Hu et al., 2018; Ottesen et al., 2013). An alternative -omics approach that focuses more on taxonomic diversity analyses and as such reducing sequencing costs compared with full metagenomic and metatranscriptomic analyses is mitochondrial enrichment (mitogenomics) (Andújar et al., 2015; Liu et al., 2016) – an approach yet to be implemented for studying any microbial group including protists.

### Contemporary best practices for protist community analyses

Despite some drawbacks, metabarcoding of the 18S rRNA gene V4 and V9 region represents by far the most cost-efficient way to assess protist environmental diversity across large spatio-temporal scales (i.e. large number of samples). We emphasise that no true eukaryote-wide universal primer pair without biases can be designed for protists due to both the paraphyletic nature of protists and the extreme phylogenetic diversity they encompass. A cumulative overview and information of primer sets used to date that should give an overview for researchers interested in studying protists is shown in Table 1 that together with the other figures provide an overview of advantages and disadvantages. These show, for instance, that the by far most commonly used primer pair 8 (Stoeck et al., 2010) to date, has clear disadvantages and only matches about 67% of all protists (Table 1), especially prevalent in the most common soil protists: Amoebozoa and Rhizaria. Many other primer pairs that can be applied using the highest throughput Illumina sequencing approaches face similar biases and perform only slightly better (Table 1). Therefore, an unbiased comparison between environments and using data obtained so far is difficult and we cannot suggest an ideal primer pair for exhaustive protist community profiling across systems.

Ideally, long-read sequencing using PacBio or Nanopore sequencing is needed to cover most protistan diversity in a sample using primer pair 6 as it covers about 93% of all protists (Table 1). An alternative is accepting shorter read lengths and as such lower phylogenetic resolution using partial single-ended amplicon analyses (Pauvert et al., 2019) with primer pair 6. Pairing sequence reads reduces the possibilities of primers to select from as their amplicon sizes are longer than the most commonly used Illumina platforms can amplify. This means, a lower diversity of protists can be targeted using 2×300bp, 2×250bp or 2×150bp sequencing with the best primers amplifying 88% (primer pair 6), 75% (34) or 63% (29) of protistan diversity, respectively. If whole microbiome analyses including protists, bacteria, archaea and fungi are envisioned PCR-free metagenomics and metatranscriptomics are suggested, but their costs currently still commonly are preventing large-scaled analyses. A more cost-efficient alternative to get a cumulative microbiome overview is using primer pairs such as 33 that amplify both prokaryotes and eukaryotes.

We summarize the primer selection for a given study system in a user-friendly diagram in Fig. 5. This overview together with additional information provided in this manuscript should help researchers, particularly ecologists, to choose a primer pair that best suits the research question, system studied and sequence platform available. This should facilitate analyses of protistan diversity within mainstream ecological and microbiome studies, and also applied research such as biomonitoring and bioindication (Stefan Geisen et al., 2018; J. Pawlowski, Lejzerowicz, Apotheloz-Perret-Gentil, Visco, & Esling, 2016). Following this guide, protist community analyses will also become more comparable between studies leading to a greater capacity for, and more meaningful, comparisons of protist communities across eco-systems, similar to that possible for bacteria (Knight et al., 2018) and fungi (Nilsson et al., 2019). Our intention is that this guide develops over time, therefore we intend to regularly update the available metabarcoding primers to study protists (https://github.com/pr2database/pr2-primers). We ask the community for their active input in keeping this work up-to-date, in order to fully establish protist community profiling as a standard tool in future microbiome studies (https://github.com/pr2database/pr2-primers/issues).

**Fig. 5.**
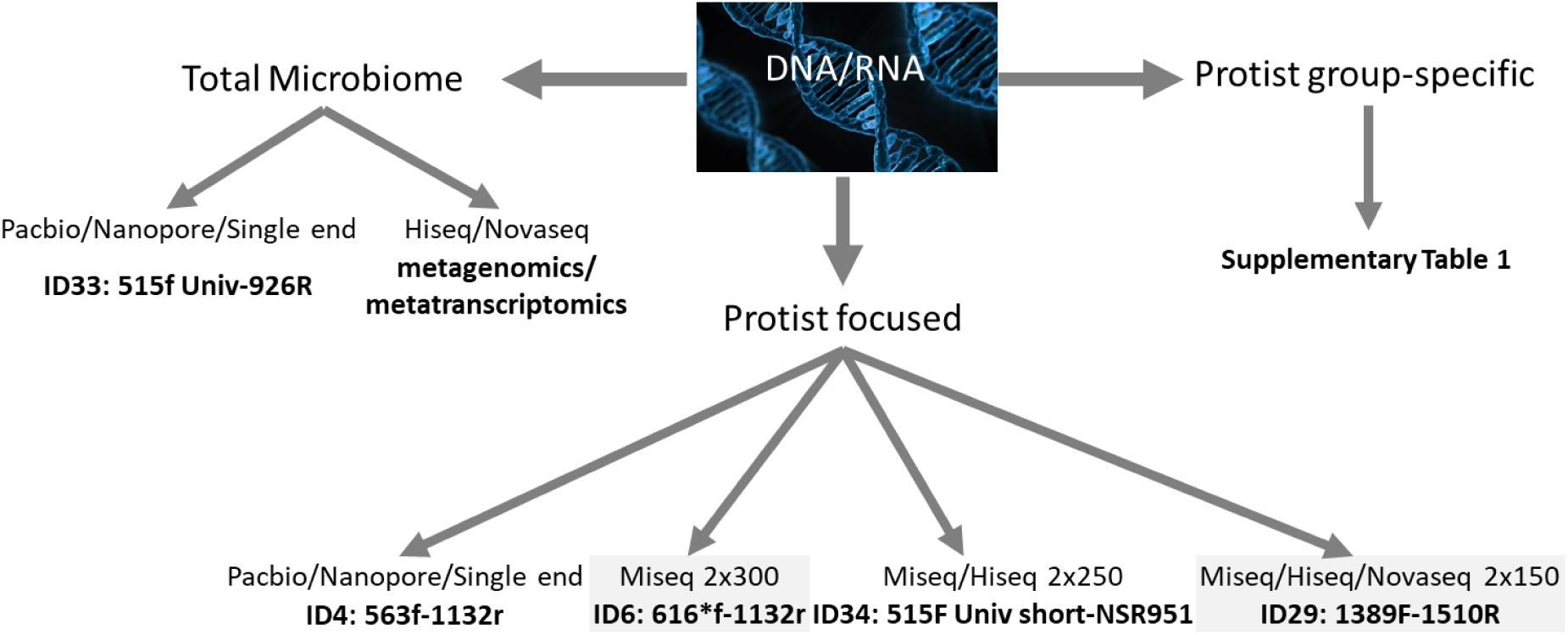
Decision-making chart to guide molecular approaches for protist community analyses. The ideal primer choice depends on the available sequencing platforms and the study question (protist- focused or microbiome- that target all prokaryotes and eukaryotes simultaneously). The longer the sequencing read, the broader theoretical coverage of protistan diversity. See Table 1 and main Figs. for detailed information on the respective primers.

## Methodology

18S rRNA gene primer sets used in metabarcoding studies were collected from literature. Primer sequences and primer sets (knowing that several primer sets may share at least one primer) were stored in a custom MySQL database (full list available at https://github.com/pr2database/pr2-primers/wiki/18S-rRNA-primer-sets and https://github.com/pr2database/pr2-primers/wiki/18S-rRNA-primers). Primer sets targeting the V4 and V9 region of the 18S rRNA gene were selected to determine in silico amplification of sequences stored in version 4.12.0 of the PR^2^ database (Guillou et al. 2013, https://github.com/pr2database/pr2database/releases/tag/v4.12.0). Sequences with ambiguities were discarded. For V4 and V9, sequences with length shorter than 1200 and 1650 bp, respectively, were not considered. Moreover, for V9, since many 18S rRNA do not cover the full V9 region, we only kept sequences that contains the canonical sequence GGATC[AT] which is located at the end of the V9 region, just before the start of the internal transcribed spacer 1. A R script was used to compute the % of sequences matching the forward, reverse and both primers using the Biostrings package function vmatchPattern() with the following parameters: max.mismatch=0, min.mismatch=0, with.indels=FALSE, fixed=FALSE, algorithm=“auto”. The data were tabulated using the dplyr package and plotted using the ggplot2 package. All scripts are available at https://github.com/pr2database/pr2-primers.

## Acknowledgements

This study was funded by the Gordon and Betty Moore Foundation through grant GBMF5257 (UniEuk) and the International Society of Protistologists. SG was supported by an NWO-VENI grant (016.Veni.181.078) from the Netherlands Organisation for Scientific Research. This work was also supported by NERC grants NE/H009426/1 (DB, CB) and NE/H000887/1 (DB), and by funding from the UK Department of Environment, Food and Rural Affairs (Defra) under contract FC1214 (to DB). EL was supported by a grant “Atracción de Talento Investigador” by the Community of Madrid (<u>2017-T1/AMB-5210</u>). CdV was supported by the French Government “Investissements d’Avenir” program OCEANOMICS (ANR-11-BTBR-0008).

We thank Linda Amaral-Zettler, Sophie Arnaud-Haond, Sandra Baldauf, Sergio Balzano, Cedric Berney Jens Boenigk, Jane Carlton, Simon Creer, Didier Debroas, Micah Dunthorn, Javier del Campo, Isabelle Domaizon, Virginia Edgcomb, Bente Edwardsen, Noah Fierer, Laure Guillou, Laura Katz, Patrick Keeling, Rob Knight, Franck Lejzerowicz, Purificacion Lopez-Garcia, Connie Lovejoy, Ramon Massana, Sebastian Metz, Edward Mitchell, Angela Oliverio, Xavier Pochon, Chris Seppey, David Singer, Alexey Smirnov, Thorsten Stoeck, Alexandra Worden, Adriana Zingone for initial discussions on primer choices for eukaryotes.

## Data Accessibility

All data used in this work are included in this manuscript. All scripts used are available at https://github.com/pr2database/pr2-primers. All data and primer information will continuously be updated at https://github.com/pr2database/pr2-primers/issues).

## Author Contribution

SG and DB designed the study; SG, DV, FM and EL drafted the tables with help from all authors; DV and SG created the figures; SG and DB wrote the first draft of the manuscript, supplemented with comments from all authors.

## Supporting Information

**Supplementary Table 1.**
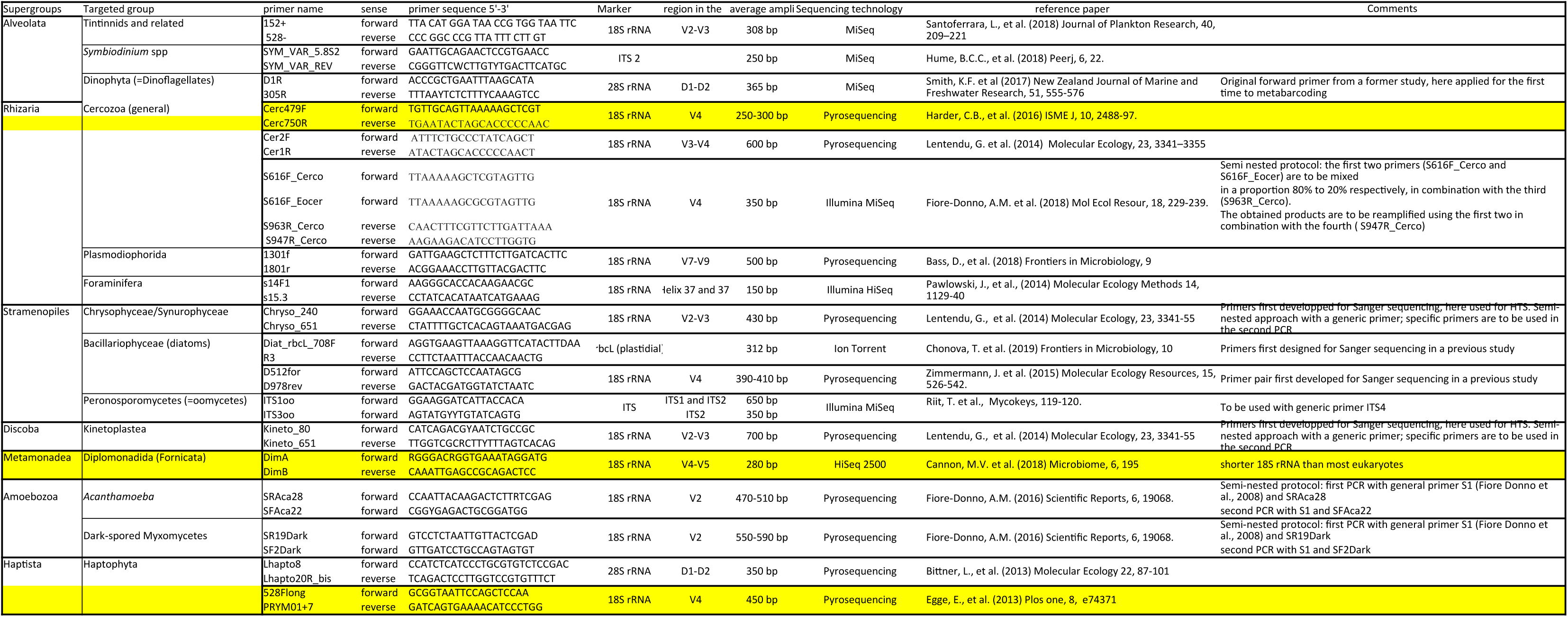
List of protist group-specific primers so far used in high-throughput sequencing based metabarcoding studies

**Suppl. Fig. 1.**
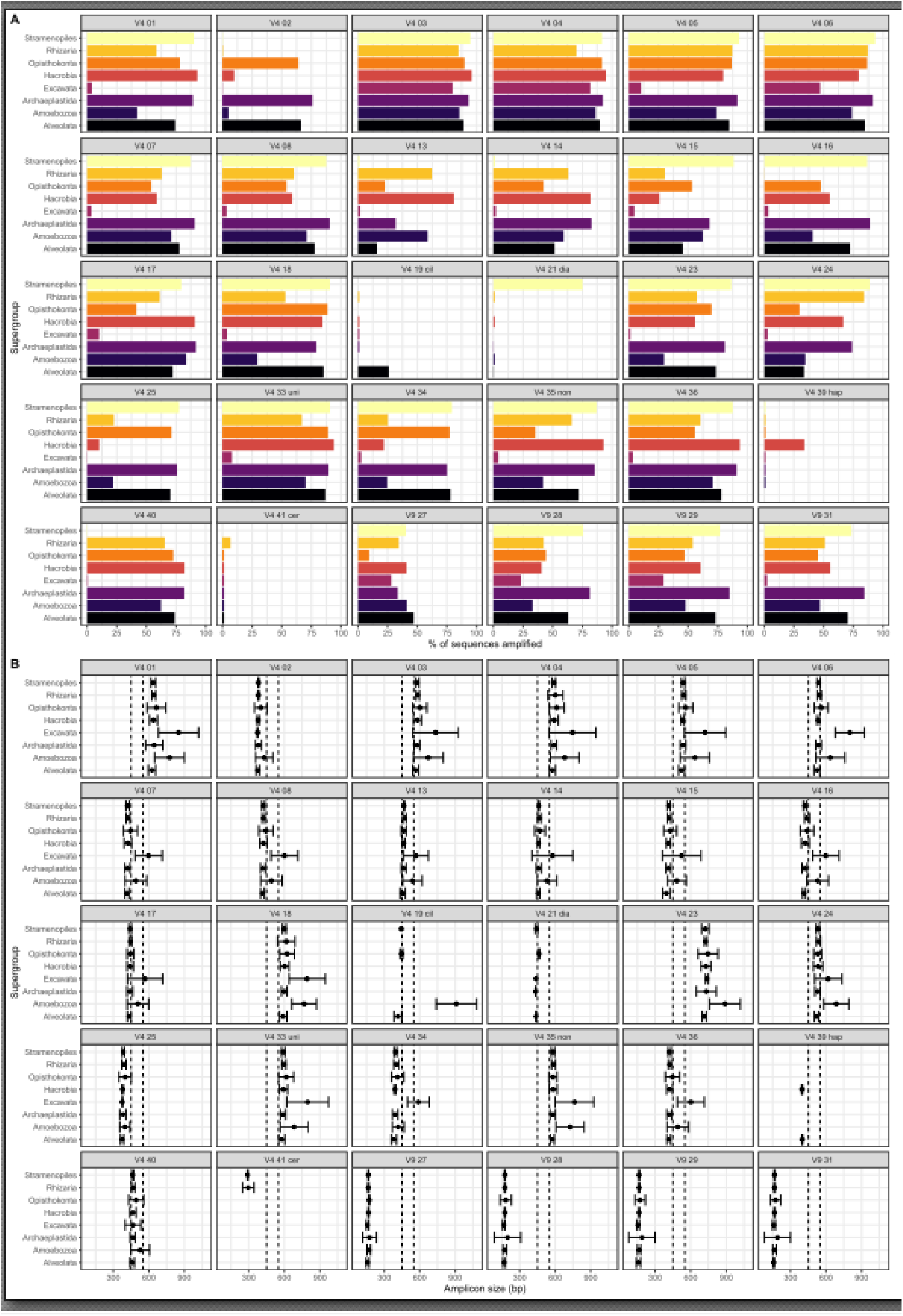
Coverage of all primer pairs used so far in high-throughput sequencing studies as identified with perfect matches of both primer pairs to the target sequences for the main protist lineages. This shows that (A) primers do not equally amplify higher taxonomic level lineages of protists and (B) amplicon lengths differ between supergroups and depending on primer sets. Note that Hacrobia represents the sum of Haptophytes, Cryptophytes and Centrohelids.

## Notes

https://github.com/pr2database/pr2-primers

